# *D. melanogaster* meiotic driver Stellate compromises sperm development by impeding a process of nuclear envelope remodeling

**DOI:** 10.1101/2025.09.29.679412

**Authors:** Xuefeng Meng, Yukiko M. Yamashita

## Abstract

Meiotic drive is a phenomenon that violates Mendel’s Law of Equal Segregation, leading to biased transmission of the meiotic driver to the offspring. *D. melanogaster* Stellate (Ste) is an X-linked meiotic driver that preferentially harms Y-chromosome-bearing spermatids, thereby favoring the transmission of the X chromosome to the next generation. We have recently shown that Ste protein segregates asymmetrically during meiosis I with a strong bias toward the Y-chromosome-inheriting side, leading to the eventual demise of the Y-chromosome-containing spermatids. However, the cellular mechanisms by which Ste protein interferes with spermatid development remain unknown. Here, we show that Ste-containing spermatids are delayed in the process of nuclear envelope remodeling, an essential process during sperm DNA compaction. We show that components of the nuclear lamina (such as Lamin Dm0, and the LEM domain proteins Otefin and Bocks) are rapidly removed during nuclear envelope remodeling during the early stages of normal spermatid development. However, Ste-containing spermatids retained these nuclear lamina proteins for a prolonged time. Their delayed removal is associated with defective formation of the dense complex, which is composed of a bundle of microtubules and serves as a structural support for sperm nuclear morphogenesis. Defective dense complex formation in Ste-containing spermatids led to defective sperm DNA compaction. Together, the present study reveals an unexpected cellular mechanism by which a meiotic driver, Ste, sabotages sperm development.

**Article summary:** Stellate is an X-chromosome-linked meiotic driver in *D. melanogaster*, which interferes with the process of spermatogenesis and causes preferential death of the Y-chromosome-containing spermatids. However, the cellular mechanisms by which Stellate interferes with spermatogenesis remain unknown. This study shows that Stellate-containing spermatids are defective in the process of nuclear envelope remodeling, an essential process during sperm DNA compaction. Defective nuclear envelope remodeling was associated with a failure to assemble the dense complex, a microtubule-rich structure that serves as structural support for sperm nuclear morphogenesis. Together, the study provides insights into a cellular strategy employed by a meiotic driver.

## Introduction

Meiotic drivers exploit the processes of meiosis and post-meiotic processes to bias their transmission to the offspring, violating Mendel’s Law of Equal Segregation. Meiotic drivers that operate in female meiosis can utilize the inherent asymmetry of the process that generates oocyte and polar body: female meiotic drivers may preferentially segregate to the oocytes, thereby achieving a rate of transmission higher than expected from Mendel’s Law of Equal Segregation (Chmatal *et al*. 2017; Searle and Pardo-Manuel de Villena 2024). Other meiotic drivers actively sabotage their opponents, where ‘poison/killer’ specifically compromises the gametogenesis process of the target (opponent) allele. Such meiotic drivers are reported broadly from fungi to mouse (Bravo Nunez *et al*. 2018; Courret *et al*. 2019; Srinivasa and Zanders 2020; Courret *et al*. 2023).

In Drosophila, a handful of male meiotic drivers that sabotage spermatogenesis (‘sperm-killing’ drivers) have been identified, including *D. melanogaster*’s *Segregation Distorter (SD)* (Sandler *et al*. 1959; Larracuente and Presgraves 2012) and *D. simulans*’ *Sex Ratio (SR)* (Tao *et al*. 2001; Montchamp-Moreau 2006; Tao *et al*. 2007a). *D. melanogaster*’s *Stellate (Ste)* has been suspected to be a meiotic driver based on a weak female-biased sex ratio, but whether it is indeed a meiotic driver has been debated for decades (Hurst 1992; Palumbo *et al*. 1994; Hurst 1996; Robbins *et al*. 1996; Belloni *et al*. 2002). We recently showed that Ste is indeed a meiotic driver that kills Y-chromosome-bearing spermatids during spermatogenesis (Meng and Yamashita 2025). Although genes involved in these meiotic drive systems do not apparently belong to a common pathway (RanGAP for *SD*, Casein kinase II beta subunit homolog for Ste, and genes containing protamine-like domains and siRNAs targeting them for *D. simulans SR*)(Bozzetti *et al*. 1995; Merrill *et al*. 1999; Muirhead and Presgraves 2021; Vedanayagam *et al*. 2021), many exhibit a cytological phenotype that is strikingly similar to each other, where the affected spermatids fail to undergo proper sperm DNA compaction, associated with defects in proper incorporation of protamine, sperm-specific DNA packaging proteins, into sperm DNA (Tao *et al*. 2007b; Courret *et al*. 2019; Herbette *et al*. 2021; Courret *et al*. 2023; Meng and Yamashita 2025). These observations may suggest the presence of a common biological process that is vulnerable to meiotic drive. However, the underlying mechanism of how spermatogenesis is sabotaged is poorly understood (Courret *et al*. 2023).

Here, we describe the cellular mechanism by which Ste protein impedes sperm development in *D. melanogaster*. We found that Ste-containing spermatids fail to properly remove components of the nuclear lamina during nuclear envelope remodeling, an essential process involved in sperm DNA compaction. Although the process of nuclear envelope remodeling has been only documented at the ultrastructural level (Tokuyasu 1974), we found that the components of nuclear lamina, such as Lamin Dm0 and LEM domain proteins (Ote and Bocks), are rapidly removed from the nuclear envelope in post-meiotic spermatids, likely reflecting the process of the nuclear envelope remodeling. We found that Ste-containing spermatids are delayed in the removal of nuclear lamina components. We further show that failed nuclear envelope remodeling is accompanied by defective formation of the dense complex (also known as dense body or dense material), the structure made of a microtubule bundle and other associated proteins that serves as a structural support for sperm nuclear morphogenesis (Tokuyasu 1974; Li *et al*. 2023). Together, this study provides insights into cellular processes that are exploited by a meiotic driver.

## Materials and Methods

### Fly husbandry and strains used

All *Drosophila melanogaster* strains were raised on standard Bloomington medium at 25 °C. The following strains were used: the standard lab wild-type strain *y w* (*y^1^w^1^*, used as *X^Ste40^*), *X^Ste200^, X^Ste200^; bam-gal4* (*X^Ste200^* is described in (Meng and Yamashita 2025)), *bam-gal4:VP16* (Bloomington Drosophila Stock Center (BDSC) 80579) (Chen and Mckearin 2003), *Mst27D-mCherry* (BDSC 95385, 95396), *Mst27D-GFP* (BDSC 95389), *mRFP*-*Mst77F* (Kyoto Stock Center DGRC 118101)(Jayaramaiah raja and Renkawitz-pohl 2005), *UAS-Lamin Dm0^RNAi^* (TRiPHMC04816, BDSC 57501), *Pros*α*2-GFP, Pros*α*6T-GFP* (Zhong and Belote 2007; Palacios *et al*. 2021), *UAS-Pros*α*4T1^RNAi^* (TRiPHMJ23606, BDSC 61980).

### Immunofluorescence staining and microscopy

Testes from 0- to 3-day-old males were dissected in 1x PBS and fixed in 4% formaldehyde in 1x PBS for 30 minutes. Fixed testes were then washed in 1x PBST (PBS containing 0.1% Triton X-100) for at least 2 hours, followed by incubation with primary antibodies diluted in 1x PBST containing 3% BSA at 4 °C overnight. Samples were washed three times in 1x PBST for 30 minutes each and then incubated with secondary antibodies in 1x PBST with 3% BSA at 4 °C overnight. After a similar washing procedure, samples were mounted in VECTASHIELD with DAPI (Vector Labs). Images were acquired using a Leica Stellaris 8 confocal microscope with a 63x oil immersion objective lens (numerical aperture 1.4) and processed with Adobe Photoshop software. The primary antibodies used were: anti-Ste (1:200; guinea pig) (Venkei *et al*. 2023), anti-Lamin Dm0, anti-Lamin C antibodies (1:200, Developmental Studies Hybridoma Bank), anti-Otefin antibody (1:100, raised in rabbit against a peptide: CEYKSKVVEPPRRQVY, LabCorp) and anti-Bocks antibody (1:100, raised in rabbit against a peptide: CPRIEPSTYRPTDLG, LabCorp). Phalloidin-Alexa Fluor 568 (1:200; Thermo Fisher Scientific, A12380) was used to stain F-actin. Alexa Fluor-conjugated secondary antibodies (Life Technologies) were used at a 1:200 dilution.

### Data availability

All data are provided in the manuscript.

## Results

### Stellate-positive spermatids are defective in nuclear envelope remodeling

Stellate (Ste) is a sperm-killing meiotic driver, impacting the process of sperm DNA compaction (Fig. 1A) (Meng and Yamashita 2025). In Drosophila, spermatids develop in synchrony within a cyst that contains a cluster of 64 nuclei resulting from 4 mitotic divisions and 2 meiotic divisions (Fuller 1993; Yamashita 2018). Ste protein segregates asymmetrically during meiotic divisions, and is preferentially inherited into Y-bearing spermatids during meiosis I (Fig. 1B) (Meng and Yamashita 2025). Ste also segregates asymmetrically during meiosis II, sparing half the Y-bearing spermatids (Fig. 1B) (Meng and Yamashita 2025). Due to asymmetric segregation of Ste protein during meiosis, each cyst contains both Ste-positive (Ste^+^) and Ste-negative (Ste^-^) spermatids (Fig. 1A). Ste protein later causes defects in sperm DNA compaction, leading to preferential death of Y-bearing spermatids (Fig. 1A) (Meng and Yamashita 2025). To understand the underlying cellular defects caused by Ste protein that lead to failure in sperm DNA compaction, we conducted detailed cytological characterization of spermatogenesis in the testis moderately expressing Ste protein, the condition that leads to meiotic drive phenotype, due to an elevated copy number of Ste repeats on the X chromosome (X^Ste200^)(Meng and Yamashita 2025).

**Figure 1:**
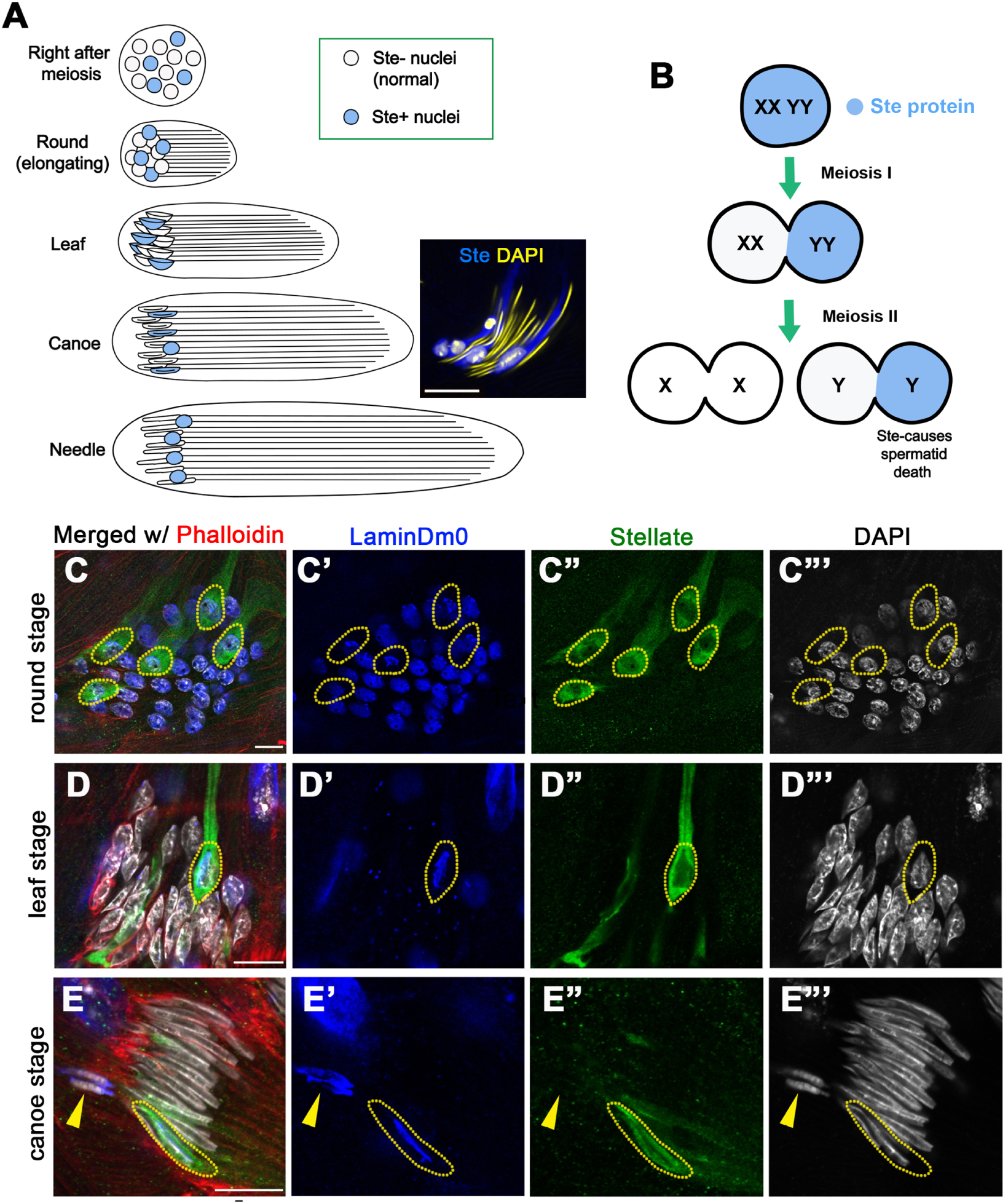
Ste^+^ spermatids are delayed in Lamin Dm0 removal during spermiogenesis. A) Schematic of spermatid development (spermiogenesis), depicting nuclear morphology changes and sperm tail elongation. 64 sperm nuclei, resulting from 4 mitotic divisions and 2 meiotic divisions, are encapsulated in a cyst and develop in synchrony. Blue nuclei indicate Ste protein-containing (Ste^+^) spermatids, which eventually fail in sperm DNA compaction. The inset shows a representative image of a cyst with Ste^+^ spermatids (Blue: Ste protein. Yellow: DAPI). B) Ste protein is expressed in spermatocytes and segregates asymmetrically during meiotic divisions. Ste protein preferentially cosegregates with Y chromosome during meiosis I. Ste protein also segregates asymmetrically during meiosis II, which rescues half of the spermatids that inherited Ste protein during meiosis I. C-E) Ste-containing cysts from X^Ste200^ animals, which derepress Ste protein moderately, stained for Ste (green), Lamin Dm0 (blue), DAPI (gray) and Phalloidin (red), through round (C), leaf (D) and canoe (E) stage spermatids. Yellow dotted lines indicate spermatid nuclei containing Ste protein. Yellow arrowhead in E indicates defective spermatids, which exhibit Lamin Dm0 staining without detectable Ste (likely because it is already degraded during late stages of spermatid death). At the round stage, n = 35/65 Ste^+^ spermatids contained Lamin Dm0, whereas n = 0/171 Ste^-^ spermatids contained Lamin Dm0. At the leaf stage, n = 19/60 Ste^+^ spermatids contained Lamin Dm0, whereas n = 0/144 Ste^-^ spermatids contained Lamin Dm0. n> 50 testes were scored. Bars: 5µm.

We previously reported that Ste^+^ spermatid nuclei that fail to compact sperm DNA are negative for protamine (such as Mst77F (Jayaramaiah Raja and Renkawitz-Pohl 2005)), which are sperm-specific DNA packaging proteins, also known as SNBP (sperm nuclear basic proteins) (Fig. S1A) (Meng and Yamashita 2025). However, upon closer examination, we found that Ste^+^ spermatids are able to initiate Mst77F incorporation in earlier stages (Fig S1B) before losing Mst77F (Fig. S1A), suggesting that protamine incorporation may not be the primary defect of Ste-mediated sperm-killing. We speculate that the loss of Mst77F from Ste^+^ defective spermatids is a consequence of defective sperm DNA compaction, rather than a cause.

While searching for potential cytological defects that are associated with Ste^+^ spermatids, we found that Ste^+^ spermatids often have more Lamin Dm0 (Lamin B of Drosophila, a component of the nuclear lamina (Gruenbaum *et al*. 1988)), compared to Ste^-^ spermatids within the same cyst (Fig 1C-E). The nuclear lamina is a fibrous meshwork structure composed of intermediate filaments (lamins), lining the inner side of the nuclear envelope and providing mechanical support for the nucleus (Lin *et al*. 2018). We found that Lamin Dm0 becomes undetectable during spermatid development in wild type: round spermatids right after meiosis exhibited Lamin Dm0 staining, but Lamin Dm0 became quickly undetectable as spermatids progress through nuclear morphogenesis (from round to leaf and canoe stages, see Figure 1A) (Fig. S2, Lamin Dm0 staining, and Fig. 1C-E, Ste^-^, normal spermatids). This likely corresponds to the process of nuclear envelope remodeling described at an ultrastructural level, during which excess nuclear envelope and nucleoplasm are eliminated (Tokuyasu 1974; Fabian and Brill 2012). Our observation that Lamin Dm0 becomes undetectable during this period suggests that nuclear envelope remodeling during spermatogenesis involves the removal of nuclear lamina, perhaps allowing the removal of excess nuclear envelope via the removal of the inner lining. The nuclear envelope remodeling results in more than 200-fold reduction in nuclear volume and tight packaging of sperm DNA by protamine. Compared to Ste^-^ normal spermatids, which had no detectable Lamin Dm0 by the leaf stage, Ste^+^ spermatids often exhibited strong Lamin Dm0 staining (Fig 1C-E, yellow dotted lines). During early stages of spermiogenesis (e.g. leaf – early canoe stage), Ste^+^ spermatid nuclei with abnormal Lamin Dm0 retention did not show clear morphological abnormalities (Fig 1D, E), suggesting that Lamin Dm0 retention precedes DNA compaction defects of Ste^+^ spermatids. We further found that other nuclear lamina components, such as LEM domain proteins Otefin and Bocks were retained in Ste^+^ spermatids (Fig. S3), confirming that an aspect of nuclear envelope remodeling is defective in Ste^+^ spermatids.

### Stellate-positive spermatids fail to properly form the dense complex during sperm nuclear transformation

We next investigated whether and how Lamin Dm0 retention of Ste^+^ spermatids may lead to sperm DNA compaction defects later, resulting in the eventual death of Ste^+^ spermatids. After meiosis, nuclei of haploid spermatids undergo a series of morphological transformations, involving changes in the overall morphology of the nucleus (from a round shape after meiosis to an eventual highly compacted ‘needle’ shape) (Fig 2A), which is accompanied by chromatin compaction supported by the replacement of histones by protamine (Fabian and Brill 2012; Rathke *et al*. 2014). It is known that the transformation of the nuclear morphology is supported by a structure called dense complex (DC)(Tokuyasu 1974; Fabian and Brill 2012; Li *et al*. 2023). The DC consists of a bundle of microtubules and is juxtaposed to the spermatid nucleus (Fig 2A, B). The DC is thought to provide a structural support for the nuclear morphology transformation, during which sperm DNA extensively compacted (Fig 2A, B) (Tokuyasu 1974; Fabian and Brill 2012; Li *et al*. 2023). In the absence of the DC, the spermatids fail to transform nuclear morphology, leading to sterility (Li *et al*. 2023).

**Figure 2:**
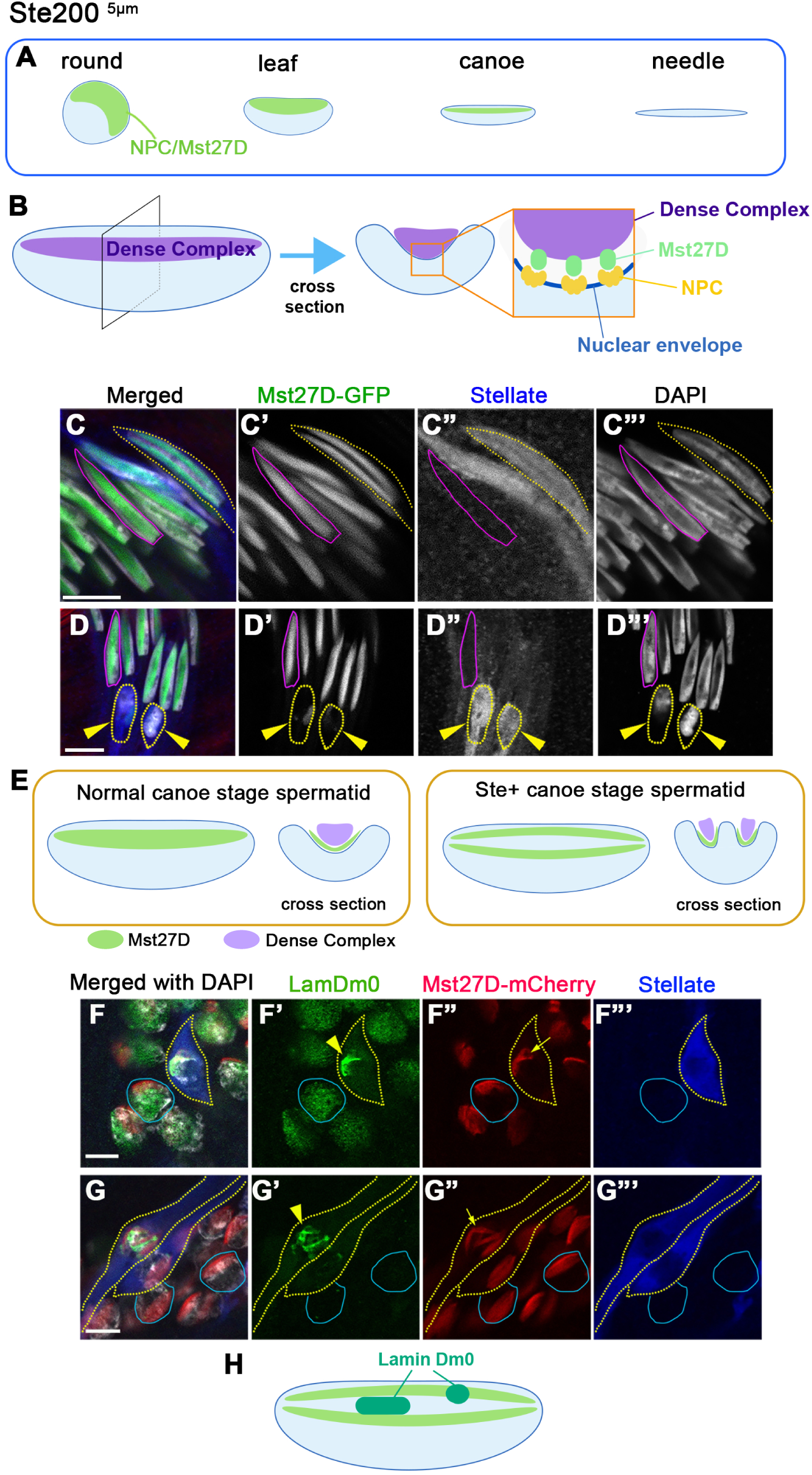
Ste^+^ spermatids are defective in dense complex formation. A) Schematic of spermatid nuclear transformation: the nuclear pore complex (NPC) polarizes to one side of the nucleus in the round stage spermatids, then localizes to the concave side of leaf-canoe stage spermatids. Upon completion of sperm DNA compaction in the needle stage, the dense complex is no longer present. Cytologically, NPCs and Mst27D, the link between NPC and the dense complex (DC), colocalize. B) Schematic of the canoe stage spermatid, depicting the nucleus and the DC, a microtubule-rich structure that serves as a structural support for sperm DNA compaction. Canoe stage spermatids exhibit a concave shape. A cross-sectional view is also shown. The dense complex is associated with the hollow of the canoe shape. Mst27D links the spermatid nuclear envelope and the DC. C-D) Immunostaining of canoe stage spermatids from X^Ste200^ animals before (C) and after (D) DNA decompaction. Both before and after DNA decompaction, Ste^+^ spermatids (yellow dotted line) exhibited an abnormal DC morphology, whereas Ste^-^ (pink line) spermatids exhibited a normal DC morphology. When sperm DNA compaction was visibly defective in Ste^+^ spermatids, the dense complex was entirely absent (D). At the canoe stage, n = 240/244 Ste^-^ spermatids had normal DC morphology, whereas n = 67/163 Ste^+^ spermatids had abnormal DC morphology, n = 10/163 Ste^+^ spermatids had no DC at all. Bars: 5µm. E) Schematic representation of the dense complex defects in Ste^+^ spermatids. F-G) Round-leaf stage spermatids from X^Ste200^ animals combined with Mst27D-mCherry (red), and stained for Ste (blue), Lamin Dm0 (green). Ste^+^ spermatids (yellow dotted lines) showed abnormal dense complex morphology, where Lamin Dm0 was located in the gap of Mst27D signal. Ste^-^ spermatids (blue lines) from the same cyst served as controls. Bars: 5µm. H) Schematic representation of Lamin Dm0 localization in Ste^+^ spermatids.

While the nuclear pore complexes (NPCs) are evenly distributed on the nuclear envelope in most cell types, including mitotic and meiotic germ cells, NPCs’ distribution becomes polarized on one side of the nuclear envelope in round spermatids (Fig 2A)(Tokuyasu 1974), a conserved feature also observed in mammalian spermatogenesis (Pereira *et al*. 2019; Manfrevola *et al*. 2021). The NPC-containing side of the nuclear envelope becomes concave, forming ‘canoe-shaped’ nucleus (Fig. 2A, B). The concave, NPC-containing side of the nuclear envelope faces the DC formed in the cytoplasm (Fig. 2B) (Tokuyasu 1974; Fabian and Brill 2012; Li *et al*. 2023). Recently, Mst27D protein was identified as a linker between NPC and DC (Fig 2B), and Mst27D was shown to be essential for the DC formation (Li *et al*. 2023). Cytologically, Mst27D and NPCs colocalize together on the NE of the developing spermatids (Fig 2A, B) (Li *et al*. 2023).

We found that DC formation was often defective in Ste^+^ spermatids. In the normal (Ste^-^) spermatids, a single DC structure forms on one side of the nucleus, visualized by the localization of Mst27D (Fig 2C, red outline)(Li *et al*. 2023). In contrast, Ste^+^ spermatids frequently had multiple DC structures that are thinner (Fig 2C, yellow dotted outline). The later Ste^+^ spermatids are often observed missing DC entirely (Fig 2D), suggesting that the defective DC that formed in Ste^+^ spermatid (Fig 2C, E) may eventually disassemble.

Co-staining of Lamin Dm0 with Mst27D-mCherry further suggested that the DC formation defects may be caused by persisting Lamin Dm0. During the round–leaf stage, when DC is being formed, Lamin Dm0 in Ste^+^ spermatid was often observed to localize to the gap of DC marked by Mst27D (Fig 2F, G), indicating that Lamin Dm0’s presence (or the nuclear envelope domain that still maintains Lamin Dm0) interferes with DC formation (Fig. 2H). Taken together, these results suggest that Ste protein may interfere with the process of NE remodeling during spermatid differentiation, leading to defective DC assembly, which in turn leads to defective sperm nuclear morphology.

### Defective removal of Lamin Dm0 contributes to sperm nuclear transformation defects in Stellate-positive spermatids

We sought to test whether the defective removal of Lamin Dm0 is causal to sperm DNA compaction defects in Ste^+^ spermatids. To this end, we introduced RNAi-mediated knockdown of Lamin Dm0 in the background of Ste-expressing X chromosome (*X^Ste200^; bam-gal4>UAS-Lamin Dm0^RNAi^*).

Ste^+^ cells are often positive for Lamin Dm0, and exhibit sperm DNA compaction defects as described above (Fig 3A). RNAi-mediated knockdown effectively depleted Lamin Dm0 from differentiating spermatids (Fig 3B”, see also Fig S4), and significantly reduced the frequency of DNA compaction defects caused by Ste (Fig 3B, C), suggesting that delayed removal of Lamin Dm0 contributes to the sperm DNA compaction defect observed in Ste^+^ spermatids. However, it is important to note that Lamin Dm0 RNAi did not completely rescue the defects of Ste^+^ spermatids (Fig 3C), suggesting that there are likely additional defects caused by Ste other than Lamin Dm0 removal, resulting in sperm DNA compaction defects. It is possible that other nuclear envelope components might still be retained in the Ste^+^ spermatids, impeding the nuclear envelope remodeling.

**Figure 3:**
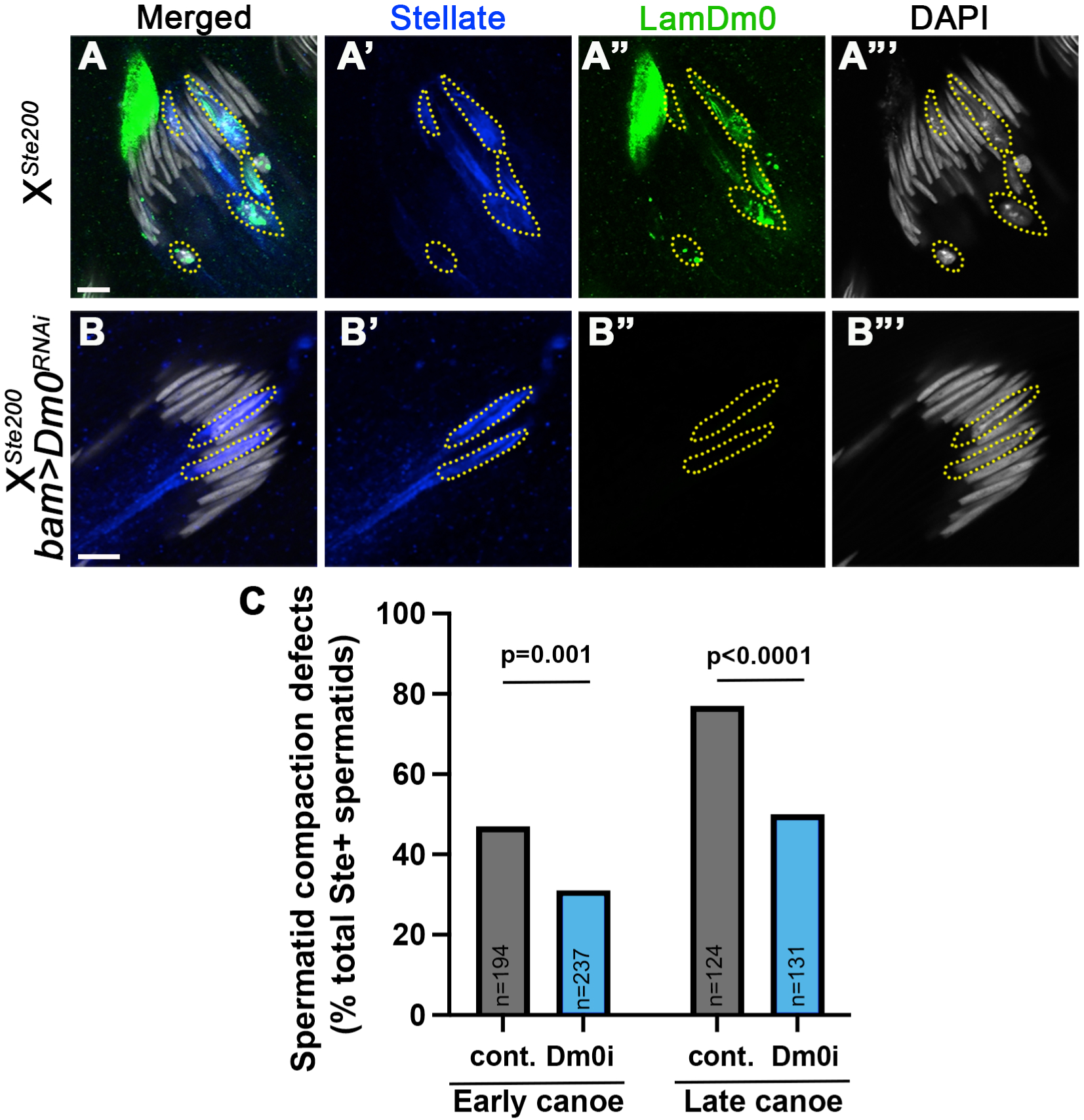
Lamin Dm0 knockdown partially rescues sperm DNA compaction defects caused by Ste. A, B) Canoe stage spermatids from X^Ste200^ animals without (A) or with (B) RNAi-mediated knockdown of Lamin Dm0 (*bam-gal4>UAS-LaminDm0^RNAi^*) stained for Ste (blue) and Lamin Dm0 (green). Yellow dotted lines indicate Ste^+^ spermatids. Bars: 5µm. C) Quantification of spermatid defects (% spermatids with defective nuclear morphology/total Ste^+^ spermatids) in indicated stages of spermatid development). N= numbers of Ste^+^ spermatids counted. More than 60 testes were scored for each genotype. P-value from the Fisher’s exact test is shown.

### The proteasome mislocalization may contribute to delayed Lamin Dm0 removal in the Ste-positive spermatids

What causes delayed removal of nuclear lamina components in Ste^+^ spermatids? We hypothesized that the proteasome pathway may play a role, because it is known that testis-specific proteasome subunits are uniquely expressed in post-meiotic spermatids and are required for spermatid development (zhong and Belote 2007). However, its substrates in the developing spermatids are not well characterized.

Interestingly, we found that the proteasome does not properly localize to the nucleus in Ste^+^ spermatids. We found that, whereas Ste^-^ spermatids exhibit nuclear localization of proteasome subunits (Prosα6T and Prosα2), as reported previously (Fig. 4) (Zhong and Belote 2007), Ste^+^ spermatids frequently lacked nuclear proteasome localization, which worsened as the spermatid differentiation progressed. Importantly, loss of nuclear proteasome in Ste^+^ spermatids became apparent from early stages of spermatid differentiation (Fig. 4A, early leaf stage) even when nuclear morphology was still normal (Fig. 4A, B). When the nuclear morphology became visibly defective, such Ste^+^ spermatids always lacked the nuclear proteasome (Fig. 4C). Interestingly, RNAi-mediated knockdown of proteasome subunit α4T1 (Prosα4T1) in germ cells (*bam-gal4>UAS-pros*α*4T1^RNAi^*) resulted in delayed removal of Lamin Dm0 in the developing spermatids. *pros*α*4T1^RNAi^* exhibited the same phenotype as reported for other proteasome subunits, with distinct sperm nuclear morphology abnormality (Fig. S5). We observed that *pros*α*4T1^RNAi^* caused delayed Lamin Dm0 removal in the round spermatid stage, when control spermatids exhibited no detectable Lamin Dm0 (Fig 5A, B). These results suggest that defective proteasome function in the nucleus of Ste^+^ spermatids may contribute to delayed Lamin Dm0 removal. However, the delay in Lamin Dm0 removal did not last as long as in Ste^+^ spermatids, and the canoe stage *pros*α*4T1^RNAi^* spermatids did not exhibit any defects in DC formation (Fig. 5C, D). These results suggest that the proteasome may contribute to, but not be essential for, Lamin Dm0 removal/NE remodeling. These results also show that Ste^+^ spermatids likely have additional defects other than defective proteasome localization, leading to sperm DNA compaction defects. Additionally, Ste^+^ spermatids do not exhibit the same phenotype as proteasome mutants, which exhibit a unique sperm nuclear morphology defect (Fig S5), which is distinct from the nuclear morphology defect of Ste spermatids. Thus, the proteasome likely has other functions beyond what is controlled by Ste to lead to its phenotype. The proteasome’s function in the cytoplasm may contribute to the phenotypes unique to proteasome mutants. In summary, defective nuclear localization of the proteasome in Ste^+^ spermatids may explain, in part, delayed Lamin Dm0 removal, although the causal relationship remains to be determined. In addition, the direct target(s) of Ste protein responsible for spermatid DNA compaction defects require further investigation.

**Figure 4:**
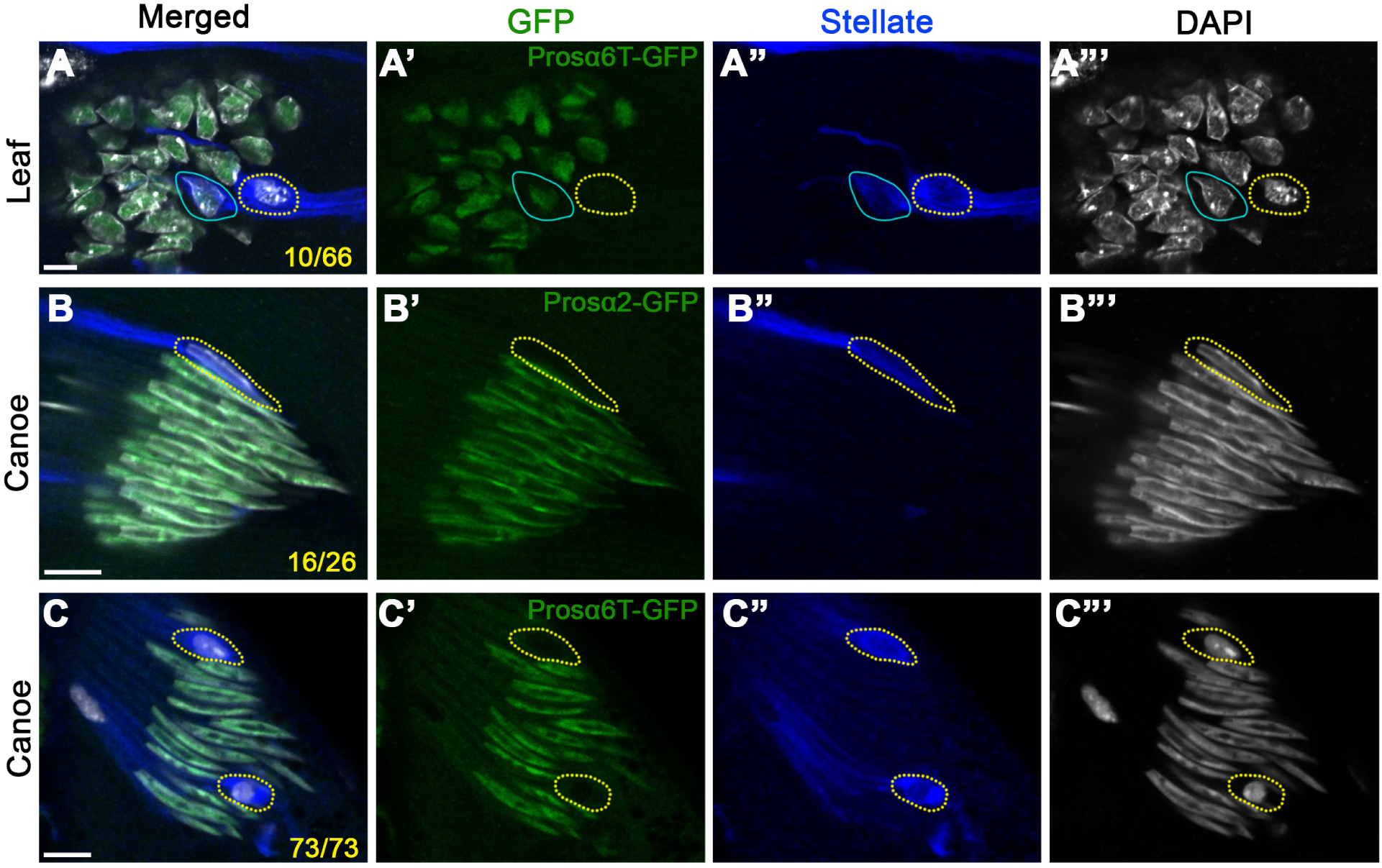
Nuclear localization of proteasome is compromised in Ste^+^ spermatids. Proteasome localization in leaf (A), canoe (B, C) stages of spermatids. Ste^+^ spermatids are outlined by yellow dotted line. Note that lack of nuclear proteasome was obvious before nuclear morphology became defective. Proteasome subunits (Prosα6T-GFP or Prosα2-GFP, Green), Ste (blue) and DAPI (gray). Bars: 5µm. Numbers in A, B, C indicate the frequency of Ste^+^ spermatids lacking nuclear proteasome.

**Figure 5:**
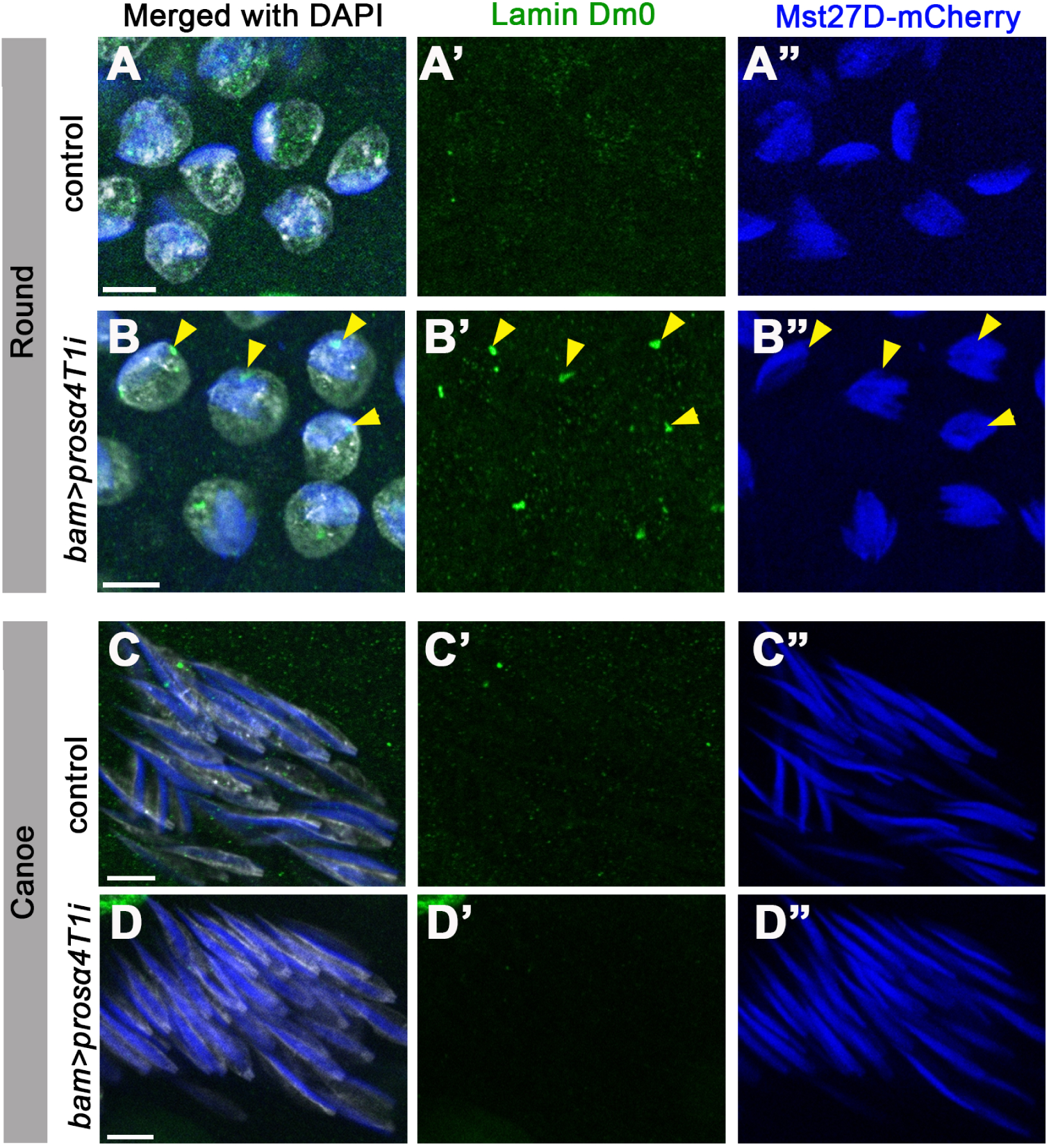
Proteasome depletion delays Lamin Dm0 removal. Round stage (A, B) and Canoe stage (C, D) spermatids stained for Lamin Dm0 (green), with the DC visualized by Mst27D-mCherry (blue). *bam-gal4>UAS-pros*α*4T1^RNAi^*, *Mst27D-mCherry* vs. sibling control lacking *bam-gal4* driver were used. In B, Lamin Dm0 foci colocalizing with Mst27D are indicated by yellow arrowheads (B). More than 40 testes were scored for each genotype. Bars: 5µm.

## Discussion

This study shows that Ste protein interferes with spermatid differentiation by targeting a rather unexpected process of spermatogenesis --- affecting the removal of nuclear lamina proteins, which likely reflects the process of nuclear envelope remodeling, previously observed by electron microscopy (Tokuyasu 1974). Although our previous study reported the lack of protamine incorporation in the Ste^+^ defective spermatid nuclei (Meng and Yamashita 2025), the present study identified a Ste-caused defect that is upstream of protamine incorporation. Our study suggests that Ste interferes with the processes of nuclear envelope remodeling, leading to defective DC formation followed by defective sperm DNA compaction.

We identified delayed Lamin Dm0 removal as a Ste-caused defect upstream of the eventual defect in protamine incorporation and sperm death. We show that Lamin Dm0 is normally removed during spermatid development: to our knowledge, this is the first report to describe the molecular signature of nuclear envelope remodeling observed by electron microscopy (Tokuyasu 1974). We found that the removal of nuclear lamina components was delayed in Ste^+^ spermatids, and our observations suggest that the retained Lamin Dm0 impedes DC formation. The fact that RNAi-mediated knockdown of Lamin Dm0 partially rescues the sperm DNA compaction defect of Ste^+^ spermatids indicates that the presence of Lamin Dm0 is a cause of defective DC formation. The DC is required for sperm nuclear morphogenesis, and has been proposed to promote the elongation of the nucleus by providing the structural support (Tokuyasu 1974; Li *et al*. 2023). Thus, defective DC formation explains why Ste^+^ spermatids fail to transform their nuclear morphology.

It is interesting to note that many sperm-killing meiotic drivers identified in Drosophila species (such as *D. melanogaster SD*, *D. simulans SR*) exhibit a very similar cytological phenotype, where affected spermatids fail to undergo sperm DNA compaction within a cyst, while unaffected nuclei proceed with sperm DNA compaction normally (Fig. 1A, inset). These results imply that the process of sperm DNA compaction is sensitive to perturbation and may be an ‘easy target’ for meiotic drivers (Courret *et al*. 2023). However, a meiotic driver must fulfill another requirement – specificity. How meiotic drivers sabotage the process specifically for its opponent, without impacting themselves, is poorly understood (Courret *et al*. 2023). In the case of Ste, preferential killing of Y-bearing spermatid is achieved by asymmetric segregation of Ste protein to the side that inherits the Y chromosome during meiosis I (Meng and Yamashita 2025). This suggests that Ste might function as a ‘time bomb’ --- if Ste is specifically targeting the process of Lamin Dm0 removal, Ste protein is safe for spermatocytes, as Lamin Dm0 removal only occurs in post-meiotic spermatids. Accordingly, the spermatocytes, where Ste protein is initially produced, will not be harmed. Only after meiosis, after X and Y chromosomes are separated, Ste’s activity to interfere with the Lamin Dm0 removal becomes consequential, sabotaging the spermatids that inherited Ste during meiosis. This mechanism --- asymmetric segregation of the ‘time bomb’ (i.e. the agents that sabotage the process of spermatogenesis only at a later point of development) --- may be a powerful and generalizable strategy for meiotic drivers. This adds to the known framework of meiotic drivers, such as poison/killer specifically targeting the opponents, while utilizing antidotes to protect themselves (Srinivasa and Zanders 2020).

Taken together, this study shows the cellular mechanism by which Ste impedes sperm development. It awaits future investigation whether other meiotic drivers may employ similar strategies in achieving meiotic drive.

## Acknowledgements

We thank the members of the Yamashita lab for discussions and comments on the manuscript and help with experiments. We thank Drs. John Belote and Michael Buszczak for providing Drosophila strains. We thank the Flybase, Bloomington Stock Center, Developmental Studies Hybridoma Bank, and Kyoto Stock Center for reagents and critical information.

## References

Belloni, M., P. Tritto, M. P. Bozzetti, G. Palumbo and L. G. Robbins, 2002 Does Stellate cause meiotic drive in Drosophila melanogaster? Genetics 161: 1551–1559.

Bozzetti, M. P., S. Massari, P. Finelli, F. Meggio, L. A. Pinna et al., 1995 The Ste locus, a component of the parasitic cry-Ste system of Drosophila melanogaster, encodes a protein that forms crystals in primary spermatocytes and mimics properties of the beta subunit of casein kinase 2. Proc Natl Acad Sci U S A 92: 6067–6071.

Bravo Nunez, M. A., N. L. Nuckolls and S. E. Zanders, 2018 Genetic Villains: Killer Meiotic Drivers. Trends Genet 34: 424–433.

Chen, D., and D. M. McKearin, 2003 A discrete transcriptional silencer in the bam gene determines asymmetric division of the Drosophila germline stem cell. Development 130: 1159–1170.

Chmatal, L., R. M. Schultz, B. E. Black and M. A. Lampson, 2017 Cell Biology of Cheating-Transmission of Centromeres and Other Selfish Elements Through Asymmetric Meiosis. Prog Mol Subcell Biol 56: 377–396.

Courret, C., C. H. Chang, K. H. Wei, C. Montchamp-Moreau and A. M. Larracuente, 2019 Meiotic drive mechanisms: lessons from Drosophila. Proc Biol Sci 286: 20191430.

Courret, C., X. Wei and A. M. Larracuente, 2023 New perspectives on the causes and consequences of male meiotic drive. Curr Opin Genet Dev 83: 102111.

Fabian, L., and J. A. Brill, 2012 Drosophila spermiogenesis: Big things come from little packages. Spermatogenesis 2: 197–212.

Fuller, M. T., 1993 Spermatogenesis. In The Development of Drosophila melanogaster New York: Cold Spring Harbor Laboratory Press.

Gruenbaum, Y., Y. Landesman, B. Drees, J. W. Bare, H. Saumweber et al., 1988 Drosophila nuclear lamin precursor Dm0 is translated from either of two developmentally regulated mRNA species apparently encoded by a single gene. J Cell Biol 106: 585–596.

Herbette, M., X. Wei, C. H. Chang, A. M. Larracuente, B. Loppin et al., 2021 Distinct spermiogenic phenotypes underlie sperm elimination in the Segregation Distorter meiotic drive system. PLoS Genet 17: e1009662.

Hurst, L. D., 1992 Is Stellate a relict meiotic driver? Genetics 130: 229–230.

Hurst, L. D., 1996 Further evidence consistent with Stellate’s involvement in meiotic drive. Genetics 142: 641–643.

Jayaramaiah Raja, S., and R. Renkawitz-Pohl, 2005 Replacement by Drosophila melanogaster Protamines and Mst77F of Histones during Chromatin Condensation in Late Spermatids and Role of Sesame in the Removal of These Proteins from the Male Pronucleus. Molecular and Cellular Biology 25: 6165.

Larracuente, A. M., and D. C. Presgraves, 2012 The selfish Segregation Distorter gene complex of Drosophila melanogaster. Genetics 192: 33–53.

Li, P., G. Messina and C. F. Lehner, 2023 Nuclear elongation during spermiogenesis depends on physical linkage of nuclear pore complexes to bundled microtubules by Drosophila Mst27D. PLoS Genet 19: e1010837.

Lin, C.-J., F. Hu, R. Dubruille, J. Vedanayagam, J. Wen et al., 2018 The hpRNA/RNAi Pathway Is Essential to Resolve Intragenomic Conflict in the *Drosophila* Male Germline. Developmental Cell 46: 316–326.e315.

Manfrevola, F., F. Guillou, S. Fasano, R. Pierantoni and R. Chianese, 2021 LINCking the Nuclear Envelope to Sperm Architecture. Genes (Basel) 12.

Meng, X., and Y. M. Yamashita, 2025 Intrinsically weak sex chromosome drive through sequential asymmetric meiosis. Sci Adv 11: eadv7089.

Merrill, C., L. Bayraktaroglu, A. Kusano and B. Ganetzky, 1999 Truncated RanGAP encoded by the Segregation Distorter locus of Drosophila. Science 283: 1742–1745.

Montchamp-Moreau, C., 2006 Sex-ratio meiotic drive in Drosophila simulans: cellular mechanism, candidate genes and evolution. Biochem Soc Trans 34: 562–565.

Muirhead, C. A., and D. C. Presgraves, 2021 Satellite DNA-mediated diversification of a sex-ratio meiotic drive gene family in Drosophila. Nat Ecol Evol.

Palacios, V., G. C. Kimble, T. L. Tootle and M. Buszczak, 2021 Importin-9 regulates chromosome segregation and packaging in Drosophila germ cells. J Cell Sci.

Palumbo, G., S. Bonaccorsi, L. G. Robbins and S. Pimpinelli, 1994 Genetic analysis of Stellate elements of Drosophila melanogaster. Genetics 138: 1181–1197.

Pereira, C. D., J. B. Serrano, F. Martins, E. S. O. A. B. da Cruz and S. Rebelo, 2019 Nuclear envelope dynamics during mammalian spermatogenesis: new insights on male fertility. Biol Rev Camb Philos Soc 94: 1195–1219.

Rathke, C., W. M. Baarends, S. Awe and R. Renkawitz-Pohl, 2014 Chromatin dynamics during spermiogenesis. Biochim Biophys Acta 1839: 155–168.

Robbins, L. G., G. Palumbo, S. Bonaccorsi and S. Pimpinelli, 1996 Measuring meiotic drive. Genetics 142: 645–647.

Sandler, L., Y. Hiraizumi and I. Sandler, 1959 Meiotic Drive in Natural Populations of Drosophila Melanogaster. I. the Cytogenetic Basis of Segregation-Distortion. Genetics 44: 233–250.

Searle, J. B., and F. Pardo-Manuel de Villena, 2024 Meiotic Drive and Speciation. Annu Rev Genet 58: 341–363.

Srinivasa, A. N., and S. E. Zanders, 2020 Meiotic drive. Curr Biol 30: R627–R629.

Tao, Y., L. Araripe, S. B. Kingan, Y. Ke, H. Xiao et al., 2007a A sex-ratio meiotic drive system in Drosophila simulans. II: an X-linked distorter. PLoS Biol 5: e293.

Tao, Y., D. L. Hartl and C. C. Laurie, 2001 Sex-ratio segregation distortion associated with reproductive isolation in Drosophila. Proc Natl Acad Sci U S A 98: 13183–13188.

Tao, Y., J. P. Masly, L. Araripe, Y. Ke and D. L. Hartl, 2007b A sex-ratio meiotic drive system in Drosophila simulans. I: an autosomal suppressor. PLoS Biol 5: e292.

Tokuyasu, K. T., 1974 Dynamics of spermiogenesis in Drosophila melanogaster. IV. Nuclear transformation. J Ultrastruct Res 48: 284–303.

Vedanayagam, J., C. J. Lin and E. C. Lai, 2021 Rapid evolutionary dynamics of an expanding family of meiotic drive factors and their hpRNA suppressors. Nat Ecol Evol.

Venkei, Z. G., I. Gainetdinov, A. Bagci, M. R. Starostik, C. P. Choi et al., 2023 A maternally programmed intergenerational mechanism enables male offspring to make piRNAs from Y-linked precursor RNAs in Drosophila. Nat Cell Biol 25: 1495–1505.

Yamashita, Y. M., 2018 Subcellular Specialization and Organelle Behavior in Germ Cells. Genetics 208: 19–51.

Zhong, L., and J. M. Belote, 2007 The testis-specific proteasome subunit Prosalpha6T of D. melanogaster is required for individualization and nuclear maturation during spermatogenesis. Development 134: 3517–3525.

