## supplementary figures for "*D. melanogaster* meiotic driver Stellate compromises sperm development by impeding a process of nuclear envelope remodeling"

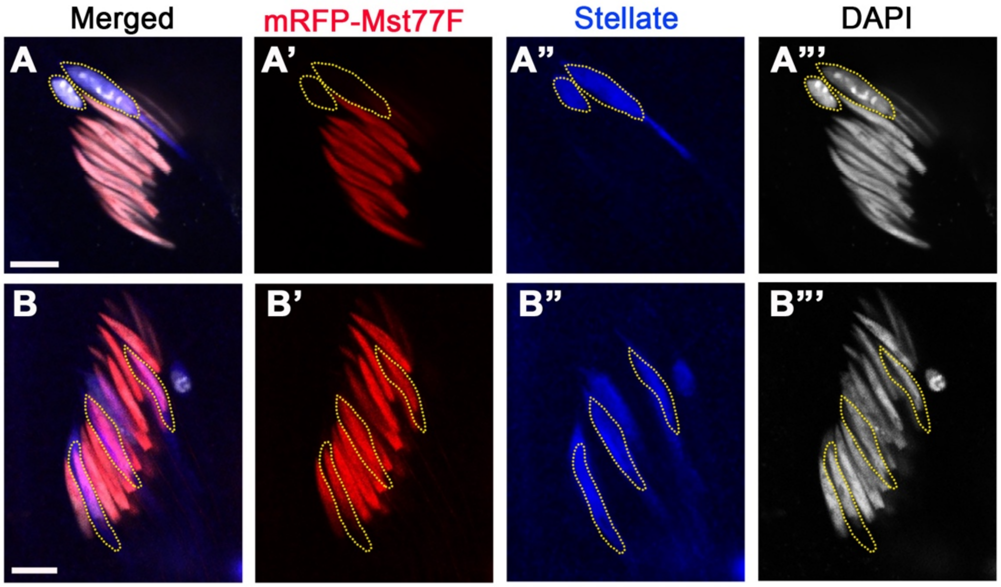


**Supplementary Figure 1. Mst77F incorporation initiates normally in Ste⁺ spermatids**

A) An example of a canoe stage spermatid cyst expressing mRFP-Mst77F, containing Ste⁺ spermatids with abnormal nuclear morphology. Stained with Ste and DAPI. When Ste⁺ nuclei exhibit abnormal morphology (yellow dotted lines), they are negative for mRFP-Mst77F.

B) An example of a canoe stage spermatid cyst expressing mRFP-Mst77F, containing Ste⁺ spermatids with normal nuclear morphology. Stained with Ste and DAPI. Ste⁺ nuclei with normal morphology (yellow dotted lines) are positive for mRFP-Mst77F.

Bars: 5µm.

30 testes (>100 canoe stage cysts) were scored.

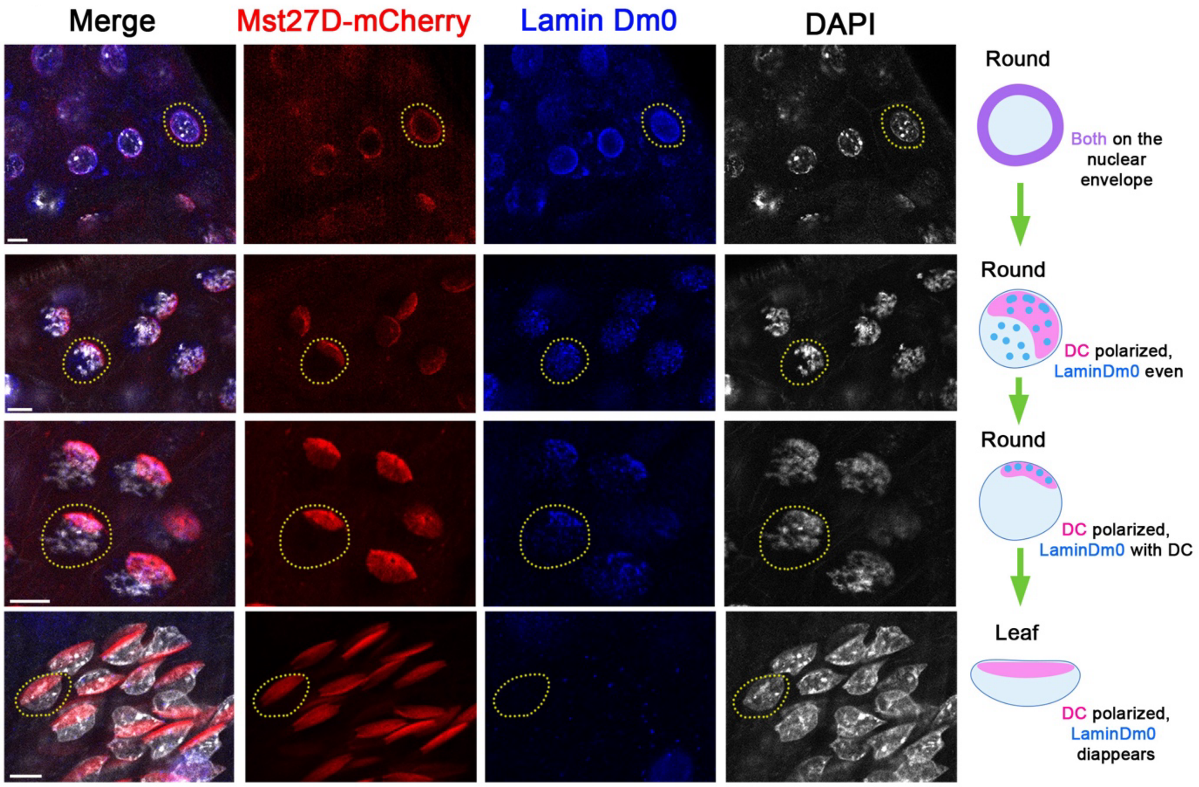


**Supplementary Figure 2. Lamin Dm0 is eliminated during spermatid nuclear morphology transformation.**

Examples of several nuclei from a cyst from round to leaf stage spermatids are shown. Mst27D-mCherry was used to visualize the localization of the DC. Stained with anti-Lamin Dm0 antibody and DAPI. Lamin Dm0, which initially localizes to the nuclear envelope, becomes progressively diminished and is completely eliminated by the leaf stage. Example nuclei are indicated by yellow dotted lines. Bars: 5µm.

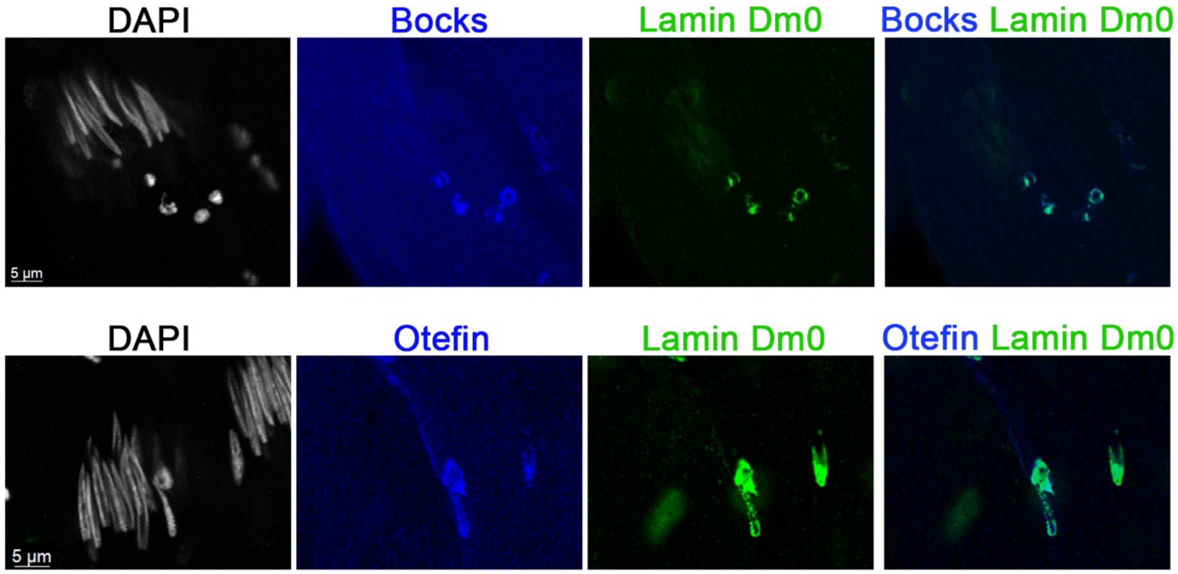


**Supplementary Figure 3. Removal of LEM domain proteins, Bocks and Otefin, is delayed in Lamin Dm0-positive spermatids**

Examples of canoe stage cyst from $X^{Ste200}$ flies stained with anti-Bocks or anti-Otefin antibodies, demonstrating that the removal of LEM domain proteins is also delayed. Bars: 5µm.

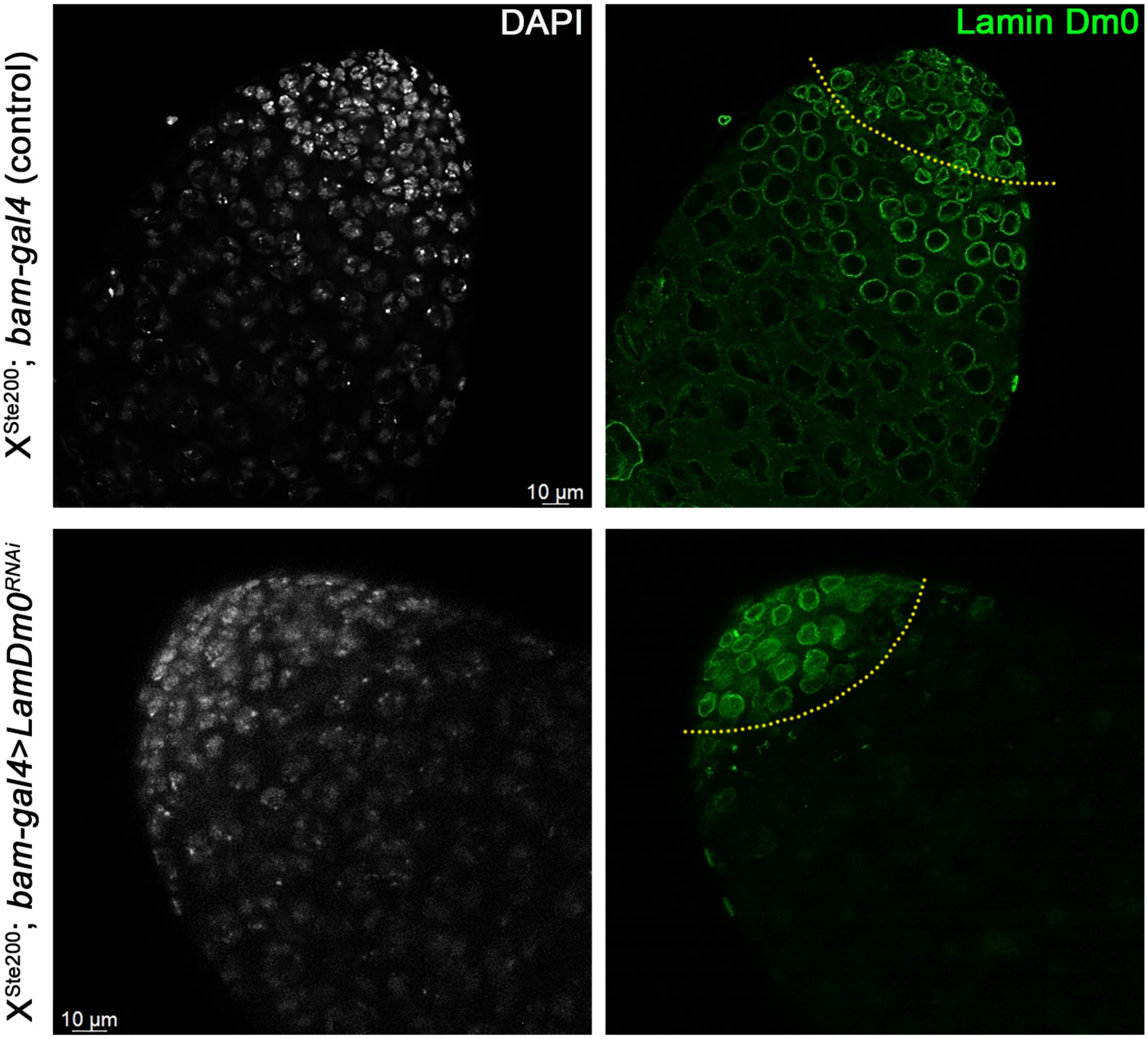


**Supplementary Figure 4. Effective knockdown of Lamin Dm0 by RNAi**

Effectiveness of *Bam-gal4>Lamin Dm0$^{RNAi}$* was confirmed by clear depletion of Lamin Dm0 staining in *bam*-expressing 4-cell spermatogonia stage and onwards (indicated by yellow dotted lines). More than 10 pairs of testes were examined for each genotype. Bars: 10 µm.

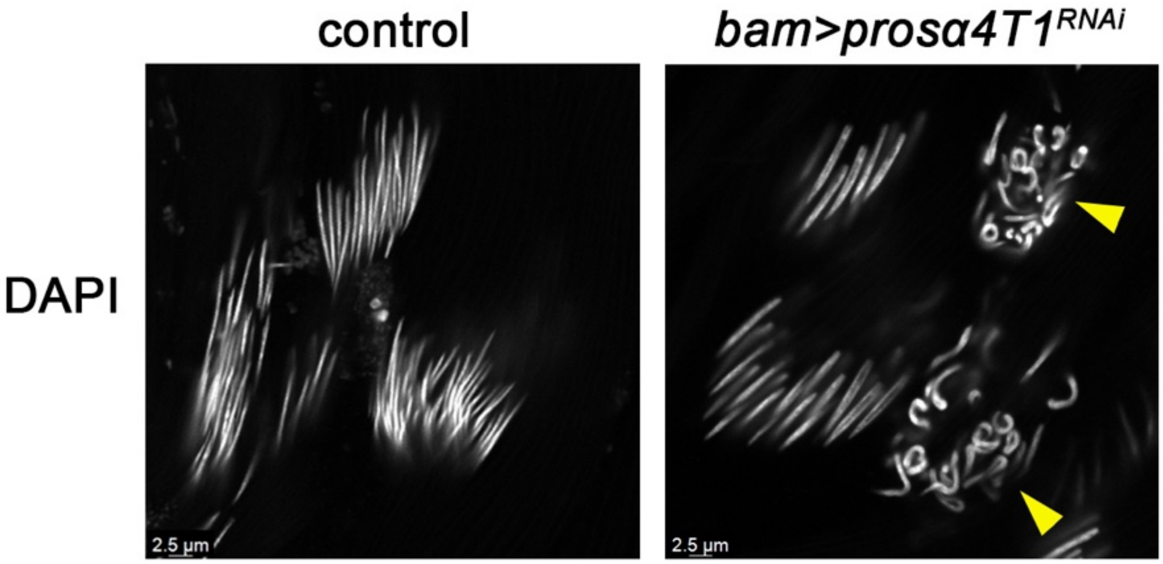


**Supplementary Figure 5.** ***pros$\alpha$4T1$^{RNAi}$*** **exhibits sperm nuclear morphology defects**
Control and *pros$\alpha$4T1$^{RNAi}$* (*bam>pros$\alpha$4T1$^{RNAi}$*) needle stage cysts stained with DAPI. N=30/30 *bam>pros$\alpha$4T1$^{RNAi}$* exhibited defective need stage nuclear morphology, equivalent to the null *pros$\alpha$6T* mutant (*pros$\alpha$6T$^{SK3}$/pros$\alpha$6T$^{SK2}$*). More than 10 testes were examined for each genotype. Bars: 2.5 µm.
